# Lateralized functional responses in the cortex arise from the dynamic interactions in the structural connectome

**DOI:** 10.1101/2020.10.16.342360

**Authors:** Neeraj Kumar, Amit Kumar Jaiswal, Dipanjan Roy, Arpan Banerjee

## Abstract

Structure-function relationships are fundamental to studies of neural systems, yet the mechanistic underpinnings of how structural networks shape hemispheric lateralization remain elusive. For instance, the asymmetric neuroanatomic embedding of primary auditory cortices was shown when connectivity with all other brain areas were considered. Concomitantly, functional lateralization e.g., left hemispheric dominance of speech processing and right hemispheric dominance of music perception, is widely acknowledged. The present article provides a parsimonious mechanistic explanation based on computational modelling and empirical recordings to demonstrate emergence of hemispheric lateralization of brain function. For validation of the model, empirical EEG recordings of auditory steady state responses (ASSR) were undertaken, and empirical findings suggest right hemispheric dominance at the level of cortical sources in binaural and monaural hearing conditions. Subsequently, we demonstrate the entrainment and phase of oscillations in connected brain regions based on a neurodynamic model constrained by empirically derived structural connectivity matrix from diffusion data. For relevance, we have taken into consideration time-delays in neural communication stemming from fibre distances and neuronal coupling arising from fibre densities. Based on relevant network parameters, we could demonstrate the emergence of right hemispheric dominance of ASSR in binaural and monaural hearing conditions when auditory cortical areas were chosen as triggers of entrained phase oscillators. Furthermore, we discovered emergence of left-lateralized functional response when bilateral Broca’s area replaced auditory areas as triggers. Hence, a single unified mechanism based on entrainment of phase oscillators in a large-scale brain network could explain both emergence of right and left hemispheric laterality.

**Significance statement:** The origin of hemispheric specialization of sensory processing is a fundamental question in neuroscience. For instance, speech and language are predominantly processed in the left hemispheric regions, while the right hemisphere is specialized for processing rhythmic, tonal, and melodic stimuli. Identification of the network mechanisms that give rise to such functional lateralization from structural constraints remains elusive. In the present study, we simulate neural activity observed during human EEG recordings of auditory steady-state responses from a biophysically realistic large-scale model constrained by underlying structural connectivity. Subsequently, we demonstrate how hemispheric lateralization of brain responses to sensory stimuli emerge from the time-delayed interactions among whole-brain neuronal ensembles.

## Introduction

Highly lateralized functional responses in the human brain have intrigued neuroscientists for a long time (Toga et al., 2003) and have tremendous clinical significance (Hecht et al., 2010). For instance, speech and language are predominantly processed in the left hemisphere (Riès et al., 2016), while the right hemisphere is specialized for processing music or rhythmic stimuli (Zatorre et al., 2001; Ross et al., 2005, Albouy et al.,2020). Depression patients have been reported to have hyper-active right hemispheres (Hecht et al., 2010). One simplistic explanation behind such observations can be attributed to the structure-function relationships in biological systems. Accordingly, attempts were made to understand the symmetries of auditory cortex embedding in the whole-brain structural connectome (Mišić et al., 2018). However, a mechanistic and causal basis of the functional asymmetry emerging from the dynamical interactions among brain areas driven by the neuro-physiological factors like conduction delays or fiber densities is poorly understood.

The present article uses computational modeling to test whether functional brain lateralization can emerge from the dynamical interactions in the whole-brain structural connectome. First, we demonstrate that auditory steady-state responses (ASSRs) from high-density EEG data are right-lateralized, which firmly establishes the validity of earlier reports (Ross et al., 2005). Second, we simulate the time-locked ASSRs from a large-scale neuro-dynamic model comprising biophysically realistic parameters, e.g., such as propagation time-delays and fiber thickness, extracted from diffusion-weighted magnetic resonance imaging (MRI). The dynamics of each parcellated (Desikan et al., 2006) brain area was modeled as a non-linear phase oscillator (Kuramoto, 1984) coupled amongst themselves via a connection matrix, the elements of which were determined by fiber densities. The time delays affecting the state variable – phase of oscillation - was dependent on the distribution of tract lengths (Abeysuriya et al., 2018) and parameterization of velocity of neural impulses along axonal tracts. Firstly, this approach allowed us to reveal how the entrainment of an environmental rhythm in any arbitrary frequency in principle could take place when the primary auditory areas are the recipient nodes of the external rhythm or synchronous drive. Secondly, it intuitively provided an understanding how external rhythms may synchronize with spontaneous oscillations at a characteristic frequency e.g., Alpha frequency as this activity gets routed from recipient auditory cortical nodes to rest of the cortical areas via physiological connection parameters e.g., time-delay. Further, we could capture that right-hemispheric dominance of ASSRs may emerge from the constraints imposed by structural connection topologies among distributed brain areas. Furthermore, to establish the model’s predictive validity for functional lateralization we constructed a scenario of virtual stimulation of bilateral Broca’s area by choosing them as the primary recipient nodes of external rhythmic input. Interestingly, based on this virtual stimulation we observed a left lateralization of entrainment of external rhythm and the power of the emergent neural oscillations. This finding demonstrates that our proposed model can also capture the left-lateralized speech and language related responses (Riès et al., 2016) as function of neuroanatomical embedding of correspondingly relevant brain areas in the whole-brain connectome.

Further, by exploring the detailed parameter space of a whole-brain model, we could tease out the biophysically realistic parameter regimes where the contributions of structural connectome shape functional lateralization. For example, the stabilization of evoked ASSRs realized as enhanced power and shifting of resonant frequency from 38-45 Hz was observed for 19-45 years old and was speculatively linked to experience-driven myelination (Poulsen et al., 2007). Our model demonstrates how a decrease in delay and increased global coupling, both factors of experience-dependent myelination, can directly enhance laterality index in young to middle-aged adults; however, middle-age to old transformation may increase time-delay due to de-myelination (Goossens et al., 2016) and, hence a decrease of laterality indices. Thus, by constructing a phenomenological model that captures hemispheric lateralization during entrainment of external rhythms in the brain, our study elucidates the dynamical principles that generate lateralization of brain function.

## Materials and methods

### Participants

Twenty-one healthy, right-handed human volunteers (16 males, 5 females, age range 22-39 years old; mean ± SD = 28 ± 2.10) participated in this study^1^. All the volunteers reported no medical history of audiological, neurological, or psychiatric disorders. All of them had normal or corrected to normal visual acuity. Informed consent were given by all the volunteers in a format approved by the Institutional Human Ethics Committee (IHEC) of National Brain Research Centre. All participants were fluent in at least two languages, Hindi and English, but some were familiar with more languages of Indian origin. All volunteers were undergraduate degree holders.

### Experimental design

Stimuli consisted of sinusoidal tones of 1 kHz frequency with a 5% rise and fall, presented 40 times per second (Linden et al., 1987). Each trial comprised of 1s “On” block (auditory stimulation) period followed by 1s “Off” block (silent) period (Figure 1A). A total of 100 trials (“On” blocks) were presented for each kind of auditory stimulation, monaural and binaural. In total, four experimental conditions, each lasting 200 seconds, were performed in the following order: 1) a baseline condition in which the volunteers were not given any tonal stimuli; 2) Binaural (in both ears); 3) Monaural left (only through left ear); 4) Monaural right (only through right ear). The time interval between each condition was set to 100 s (silent). Auditory stimuli were created and presented in Stim2 software (Compumedics, Inc., USA) at 90 dB. Participants were instructed to stay still in the sitting position, fixate on a visual cross displayed on a computer screen for the duration, and listen to the tones. When the volunteers were performing the experiment, continuous scalp EEG was recorded with relevant trigger data.

**Figure 1:**
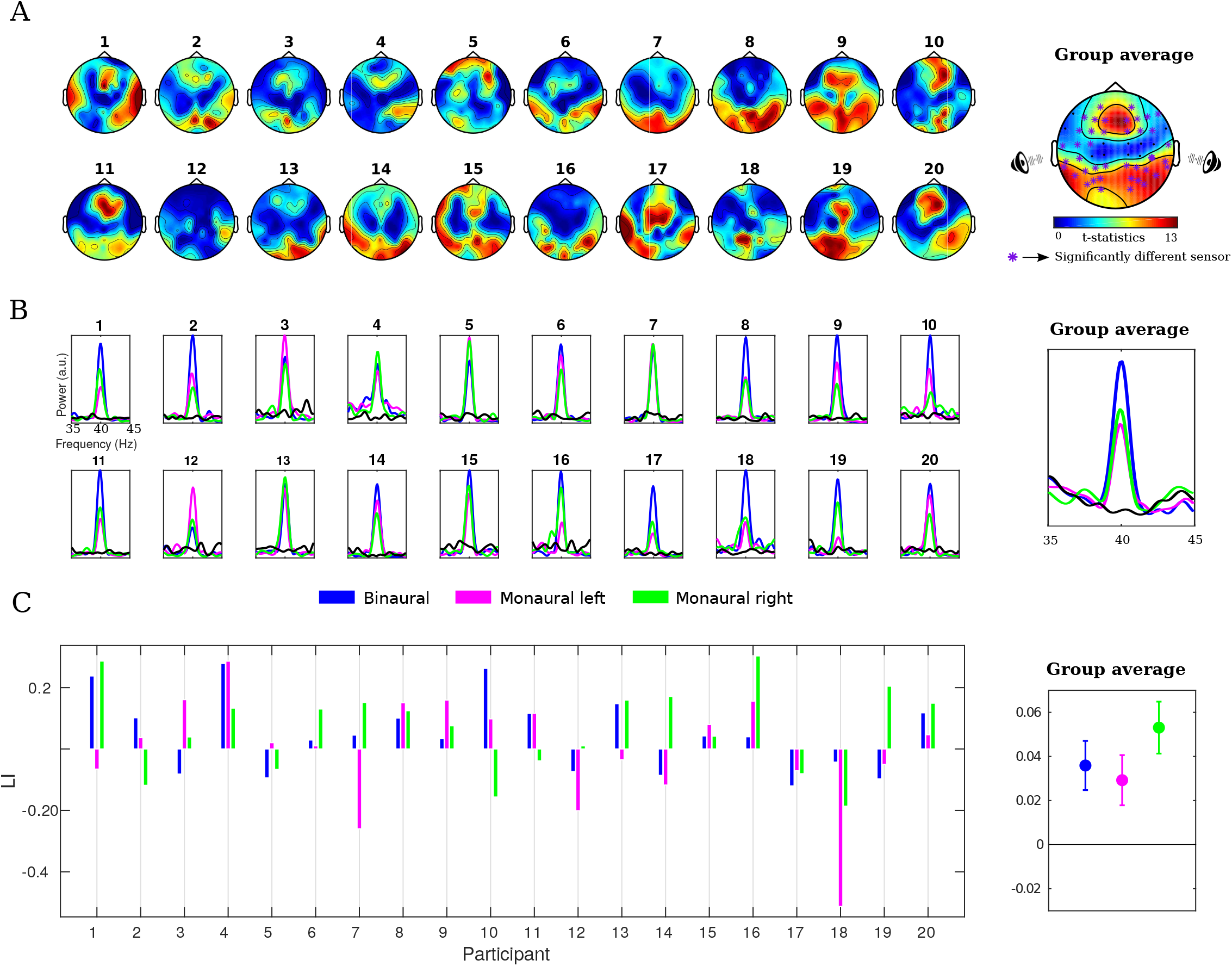
Subject-wise and group brain response. A. Topographical distribution of 40 Hz spectral power during presentation of periodic auditory stimuli at 40 Hz. The purple “*” on the topoplot maps the position of electrodes that are significantly different from the baseline condition at 95% confidence level. B. Power spectrum averaged over all sensors, measured for the monaural left (magenta), monaural right (green), binaural stimuli (blue), and baseline (black) conditions. Subject-wise responses are calculated from individual participant’s event-related potentials. For better visualization purpose, time-series was band-pass filtered in the frequency range of 35 – 45 Hz. C. Laterality indices (LIs) for 40 Hz spectral power for different stimulus conditions. The colored circle in the group plot represents the mean of LIs. Lower and upper whisker marks the lower and upper limit of 95% confidence interval, respectively.

### Neuroimaging procedure

EEG data were recorded using 64 high-density Ag/AgCl sintered electrodes mounted in an elastic head cap according to the international 10-20 system. All recordings were done in a noise-proof isolated room using NeuroScan (SynAmps2) system (CompumedicsInc, USA) with 1 kHz sampling rate. Abrasive electrolyte gel (EASYCAP) was used to make contact between EEG sensors and scalp surface, and impedance was maintained at values less than 5 kΩ in each sensor throughout the entire experiment. The EEG system-assigned reference electrode at the vertex was selected for reference, and the forehead (AFz) electrode as the ground.

### Pre-processing of EEG signals

EEG data were imported in MATLAB using EEGLAB from Neuroscan raw files. Epochs of 1000 ms from “On” blocks were extracted from each trial. The resulting epochs were then bandpass filtered to retain frequencies in the range 35 to 45 Hz, followed by detrending (baseline correction) of data to remove any linear trends from the signal. Trials having a voltage greater than ±100 μv were considered as artifacts and therefore, discarded. All trials from both monaural conditions of participant 15 were full of artefacts; therefore we had removed data of participant 15 for further analysis. Furthermore, since the objective was to estimate the steady-state activity of the brain, trials from all participants were grouped together.

### Spectral analysis

Multi-tapered power spectral density was computed using Chronux function *mtspectrumc.m* (http://chronux.org/) and customized MATLAB (www.mathworks.com) scripts at each sensor, trial, and condition. Power spectra of the concatenated time-series were calculated in the frequency range of 2 - 48 Hz having time-bandwidth product and number of tapers both set at 1, as the input parameters. Subsequently, differences in spectral power at 40 Hz (frequency of interest) between auditory stimulation tasks and baseline conditions were statistically evaluated by means of a permutation test using *ft_freqstatistics.m*, a function of FieldTrip toolbox (www.fieldtriptoolbox.org). The paired-sample t-statistic between auditory stimulation and baseline condition was computed at each sensor. Additionally, to circumvent multiple comparison problems we clustered sensors based on their spatial adjacency (Maris et al., 2007). Therefore, neighboring electrodes (minimum = 2) having individual t-value higher than corresponding *p* <0.0001 were counted as a cluster. Afterward, cluster statistics were derived by taking the sum of t - statistics across a cluster which was compared with the null distribution of cluster statistics generated by random permutation procedure (1000 times). Subsequently, the statistical significance of the spectral difference between the two conditions was assessed using a two-tailed t-test in which the observed test statistic value of the cluster was the threshold at the 95^th^ percentile of the null distribution. *p*-values of the clusters were obtained by estimating the proportion of clusters from the null distribution that are beyond the aforesaid threshold.

### Source reconstruction

We performed exact low-resolution brain electromagnetic tomography (eLORETA) (Pascual-Marqui, 2007) to locate the stimulus-specific sources of 40 Hz ASSRs. First, as a head model, we used the standardized boundary element method (BEM) volume conduction model of the human head as a common template for all participants (Oostenveld et al., 2003). We discretized the brain volume into 2807, regularly spaced three-dimensional cubic. The center of the cubic grid had the coordinates following the Montreal Neurological Institute (MNI) template. Furthermore, employing the standardized (10-20 system aligned with “colin27” brain) sensors location information along with the head model, we created a leadfield for each grid.

Subsequently, to obtain the oscillatory sources of 40 Hz activity, we employ distributed source modeling using exact low-resolution brain electromagnetic tomography (eLORETA). eLORETA estimates the current source density across brain volume by minimizing the surface Laplacian component during the construction of the spatial filter (Pascual-Marqui, 2007; Pascual-Marqui, et al., 2011). Additionally, eLORETA does not rely upon any assumption regarding the number of underlying sources while having excellent control over suppression of false positives during the detection of sources (Halder et al., 2019). The source analysis was performed using FieldTrip toolbox (Oostenveld et al., 2011; http://fieldtriptoolbox.org). The ingredients to construct a frequency domain eLORETA filter are the forward model and the cross-spectral matrix of sensor data. Hence, we computed a sensor-level cross-spectral matrix from ‘On’ blocks time series (i.e., 1 s) after re-referencing the EEG data on common average reference for all conditions. After that, a common spatial filter was computed employing combined data from all conditions. A common filter attenuates filter-specific variability during inverse modeling, i.e., the observed difference between different conditions is attributed only to the differences in conditions, not due to differences in the spatial filter. The spatial filter for each grid was then calculated in 3 orthogonal directions. Since we do not have any prior assumption about the orientation of the underlying source, the cross-spectra of sensor data were projected through the strongest orientation of dipole, i.e., denoting maximum variance. Consequently, a 3D distribution of source power across brain volume was obtained. Afterward, prominent sources were selected after thresholding the source power distribution at the 95^th^ quantile. Supra-thresholded sources were visualized by rendering onto the Colin27 brain template.

### Neurodynamic model

Structural connectivity (SC) and fiber length matrices were obtained from an online dataset derived from diffusion MRI probabilistic tractography of 40 participants of Human Connectome Project (Glasser et al., 2013; Sotiropoulos et al., 2013; Van Essen et al., 2013, Abeysuriya et al., 2018). The SC matrix and fiber length matrix were both symmetric matrices representing mean white matter densities and distance among nodes respectively. SC matrix was parcellated in 68 regions, according to the Desikan–Killiany brain atlas (Desikan et al., 2006). Subsequently, a network of 68 coupled Kuramoto oscillators was simulated, each oscillator representative of a brain parcel. The oscillators were coupled according to a coupling matrix derived from scaling and normalizing SC. Each hemisphere constituted an equal set of 34 nodes. The coupling strength matrix was normalized between 0 and 1 such that the maximum strength among connections was 1. The values at the diagonals of coupling and fiber length matrix representing self-connectivity and length with self, respectively, were set at zero.

To imbibe the realism that primary generators of ASSRs lie in auditory cortex, we chose two nodes in the auditory cortex (AC) (and other 66 nodes in the rest of the brain as non-auditory nodes (non-AC). Using the Kuramoto model (Kuramoto, 1984), the phase (θ) dynamics of any non-AC nodes is defined as,

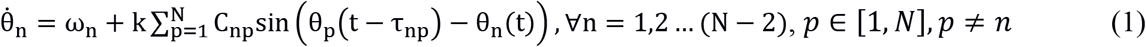

where ω_n_ represents the intrinsic frequency of the oscillator as ω_n_ = 2πf_n_; k is the mean coupling strength used to scale the all the coupling strengths; τ_np_ represents transmission delay for the propagation of information between two nodes, given the length of the fiber. Thus, 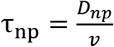 where, bio-physiologically realistic communication speed (*v*) is 5 – 20 *m/s* in adult primate brain (Ghosh et al., 2008). In equation (1) the value of p iterates from 1 to 68 including auditory nodes. This implies that phase dynamics of non-AC will depend on phase dynamics of all other nodes including both auditory nodes. However, phase dynamics of the AC nodes are defined as,

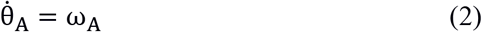

Where ω_A_ = 2π * f_A_. After setting the frequency of non-AC nodes (f_n_) at 10 Hz and frequency of AC (f_A_) nodes at 40 Hz, we investigated hemispheric laterality indices across a model parameter space, wherein k values range from 1 to 50 and v values from 5 *to* 19.1 *m/s*. During simulation for every v, time delay distribution involved in transmitting information among the network nodes was computed (Cabral et al., 2011). Similarly, structural connectivity was transformed into the coupling strength matrix C. Thus, C_np_ and τ_np_ respectively represents the coupling strength and time delay between node n and p. Therefore, the dynamics of phase (θ) at any node will be a function of its anatomical strength and distance with other nodes. The model was simulated for about 25 seconds. We took sine of (θ) obtained at each node which represents a simulation of neural time series at EEG source level. Subsequently, we calculated the power spectral density from 68 nodes followed by hemispheric laterality analysis of the spectral power at 40 Hz (See more in Laterality analysis section). During monaural condition simulations, we asymmetrically scaled the relative coupling strengths of both AC nodes with non-AC nodes by the ratio of spectral power we obtained empirically between left and right hemisphere auditory parcels after source reconstruction. Remaining procedure remained same as described above for binaural condition simulation.

### Laterality analysis

Hemispheric asymmetry in brain responses was quantified using laterality indices (*LI*), which is the difference between right hemisphere (RH) and left hemispheric (LH) responses normalized by the sum of response in both hemispheres.

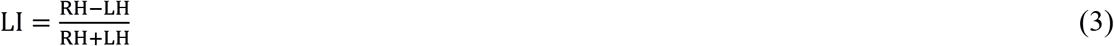

The value of *LI* lies between +1 and −1. Wherein +1 represent complete right hemispheric dominance, −1 for complete left hemispheric dominance and 0 for the bilaterally symmetric response. *LI*s for spectral power were computed at the source level for empirical and theoretical dataset.

## Results

### Spectral topography of auditory steady state responses (ASSRs)

Mean power spectra of each volunteer and the grand mean spectra across all volunteers showed enhanced spectral power, specifically at 40 Hz in both monaural and binaural conditions relative to the silent baseline (Figure 1B). Differences between the topographic scalp distribution at 40 Hz during auditory stimulations and the baseline were evaluated using cluster-based permutation tests (see Methods). Significant enhancement of spectral power at 40 Hz in distributed scalp sensor locations was observed (Figure 1A). Overall, the pattern of distribution of enhancement in the spectral power at 40 Hz was found to be similar in both binaural and monaural conditions (see extended data Figure 1-1). However, differences in magnitudes were observed across different conditions and both hemispheres. In summary, we have observed two significant clusters of spectral power during every stimulation condition. Wherein, one large cluster was located over the fronto-central area and another in bi-lateral caudal parts of the scalp (Figure 1A, extended data Figure 1-1 and Figure 1-2). However, the right posterior region channels showed greater enhancement than their counterparts in the left hemisphere during binaural and monaural right condition. The presence of right hemispheric dominance during binaural condition was cross-validated in different temporal segments (early, middle and late ERP components) of the “On” block, all three segments exhibiting right hemispheric dominance.

### Source-level functional organization of 40 Hz ASSRs

Exact low-resolution brain electromagnetic tomography (eLORETA) was used to calculate the three-dimensional spatial distribution of source activity underlying 40 Hz ASSRs. Reconstructed sources were rendered onto a standard cortical surface derived from “colin27” brain provided in the FieldTrip toolbox (http://fieldtriptoolbox.org). The locations of prominent sources during monaural left, monaural right and binaural conditions are shown in Figure 2. Anatomical labels corresponding to source regions according to the Desikan-Killiany atlas (Desikan et al., 2006) and the number of activated voxels in the respective region are summarised in Figure 2-1. Distributed sources of 40 Hz ASSRs in auditory cortices and beyond were observed. The robust bilateral activations were concentrated in the superior temporal, supramarginal, pre-central gyrus, post-central gyrus and Broca’s area. The majority of the sources exhibited right hemisphere dominance irrespective of the stimulation condition, further investigated by laterality indices (LI). Activation in the contralateral Heschl’s gyrus was observed during the monaural conditions. Left MTG showed significant activation only during monaural left and binaural conditions but not during the monaural right. Additionally, left hemisphere showed greater number of activated regions than monaural right and binaural. LI computed from source power revealed right hemispheric dominance during every type of stimulation condition, binaural (0.10), monaural left (0.13) and during monaural right (0.23) (Figure 3B).

**Figure 2:**
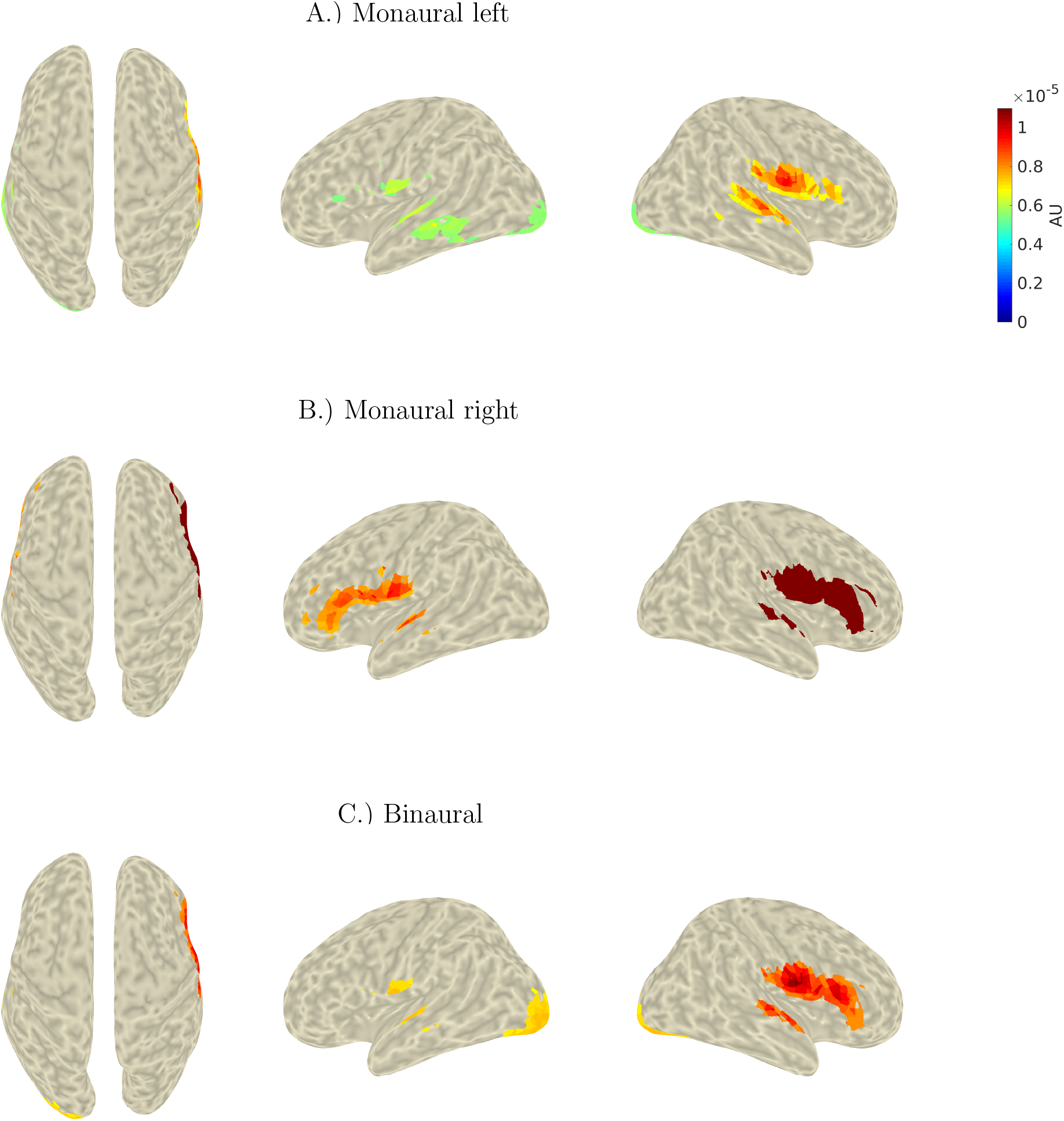
Cortical sources of 40 Hz ASSRs. Brain regions activated underlying auditory steady state responses (ASSRs) generated during 40Hz amplitude modulated stimulations for 1.) monaural left (1^st^ row), 2.) monaural right (2^nd^ row) and 3.) binaural conditions (3 ^rd^ row). Axial (1^st^ column), left (2^nd^ column) and right (3 ^rd^ column) views of brain are shown for better visualization of sources.

**Figure 3:**
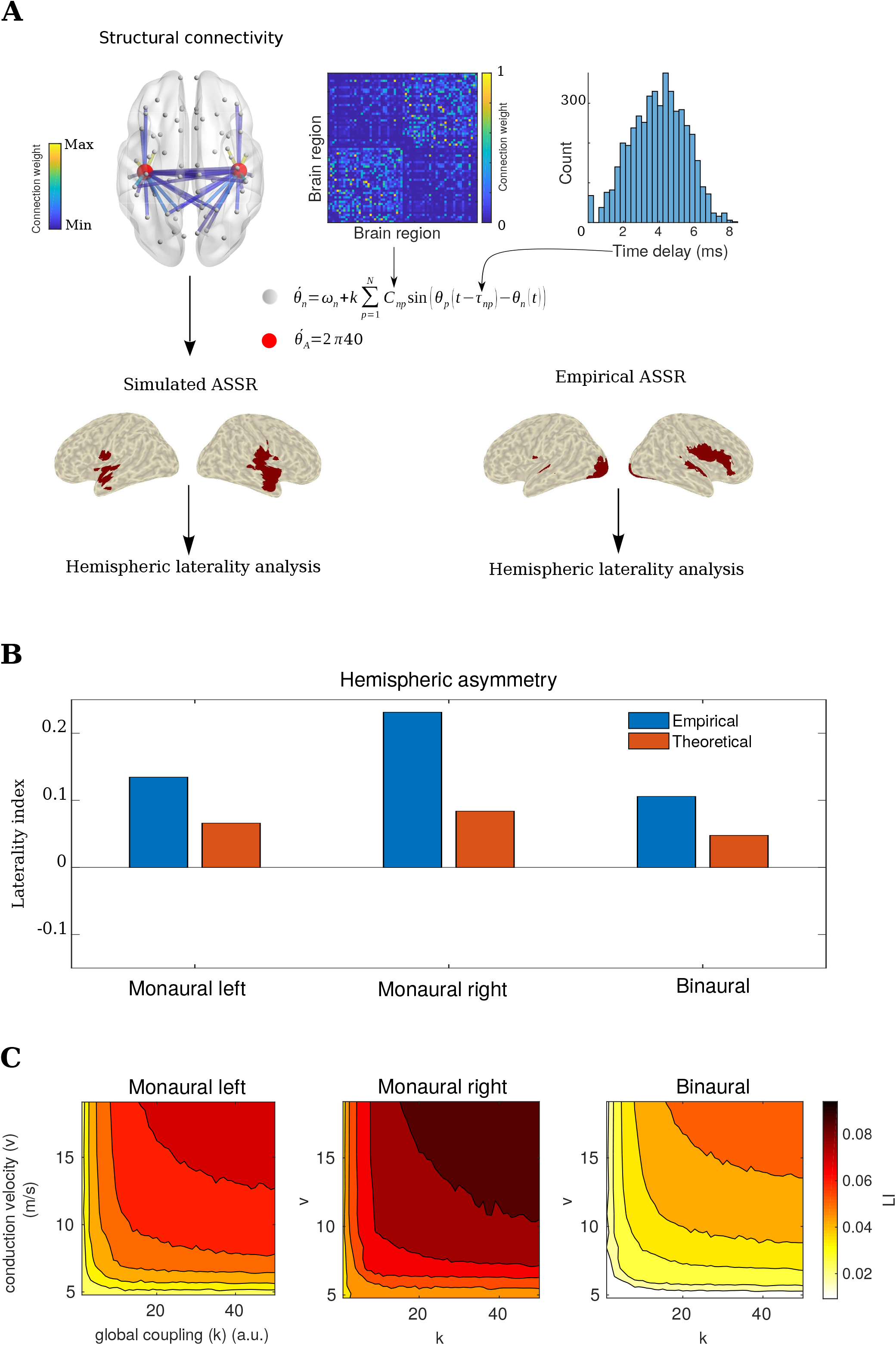
Whole-brain neuro-dynamic model to mechanistically explain ASSR lateralization. A. Pipeline to compare laterality index (*LI*) in simulated and empirical EEG data. Connection parameters of tracts among brain parcels (following Desikan-Killiany atlas) is extracted from diffusion MRI data. For ease of visualization, only the connections to auditory nodes are plotted in the glass-brain visualization. The actual model encompass the whole brain connectivity. Fiber densities guiding coupling coefficients (*C_np_*) are plotted as a matrix in the inset along with distribution of time-delay (*τ_np_*) at *v* = 19.1 *m/s*. Phase of oscillations in each parcel were modelled using Kuramoto oscillators (red circles for auditory cortical sources, grey circles for other regions, see text also). The time series obtained after reconstructing the source amplitude from phase was used for computation of laterality indices of source power at 40Hz. For illustrative purposes, results from empirical source localization analysis using eLORETA are shown for binaural condition adjacent to corresponding maps generated for simulated data. B. Bar plots of group-level mean hemispheric *LI* for 40 Hz spectral power obtained from eLORETA-based source reconstructions (Empirical) and output of neuro-dynamical model (Theoretical) C. *LI* values change under the parametric variation of global coupling (*k*) and transmission velocity (*v*) in the large-scale neuro-dynamic model.

### Neural dynamical model

Each parcellated brain area extracted from Desikan Killiany atlas (Desikan, et al., 2006) was modelled as Kuramoto phase oscillator that were coupled amongst each other. Time-delays in propagation of neural communication were considered as the time-delayed phases from rest of the brain parcels to the phase of the oscillator concerned. The time-delay and coupling coefficients that scaled the degree of influence which one brain region had over another were derived from empirical diffusion MRI probabilistic tractography data (see Methods for details, Figure 3A). The auditory cortex nodes were driven by 40 Hz external inputs while keeping other nodes’ intrinsic frequency at 10 Hz. The system of differential equations was numerically integrated using the Euler integration method for 250000-time points with a step size (*dt*) of 0.0001 representative of 25 seconds duration. Thereafter, the power spectral density was computed from the source time series of 25 seconds duration after down-sampling to 1000 Hz. The hemispheric laterality indices (*LI*) was calculated from spectral power distribution at 40 Hz. Figure 3C demonstrates the *LI* values across a range of global coupling (*k*), 1-50 and velocity of neural impulse propagation (*v*), 5 - 19.1 *m/s* (see Methods for rationale of choosing these biophysically realistic values). *k* is the overall scaling factor on individual coupling coefficients, and represent an overall strength by which all brain areas are bound as a network and has been established as a biophysically realistic parameter by earlier modelling studies (Cabral et al., 2011). We compared the *LI* obtained from the model with the empirical LI in Figure 3B for biophysically realistic *v* = 12 *m/s*. *k* = 30 and above, yielded good match with empirical LI values. The resulting LI for each condition are 0.04 (binaural), 0.06 (monaural left) and 0.08 (monaural right, Figure 3B). Positive *LI* values were obtained across the entire parameter space spanned by *k* and *v*; (0.02, 0.07] for monaural left condition, (0.03, 0.09] for monaural right condition and (0.01, 0.058] for binaural condition (Figure 3C). Since the time delay 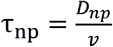 is parametrically dependent on *v, LI* is thus modulated as a function of both time delay and global coupling strength (Figure 3C). Specifically, for every type of simulation condition *LI* is increased positively (Right hemispheric dominance) when coupling (*k*) and speed of transmission (*v*) increases.

To establish the predictive validity of our hypothesis that topology of structural brain networks contributes to functional hemispheric lateralization, we constructed a scenario to explain the left hemispheric lateralization of language and speech processing. The auditory cortical nodes for entry of environmentally relevant stimuli to the whole brain connectome were replaced by parcels in bilateral Broca’s areas. According to Desikan Kiliany atlas Broca’s area encompass two parcels i.e., pars triangularis and pars opercularis. Therefore, we set the intrinsic frequency of these two parcels at 40 Hz and rest of the brain parcels at 10 Hz and numerically integrate equation 1 following steps used earlier. Corresponding *LI* values were calculated from spectral power at 40 Hz. *LI* values across the tested parameter space display two kind of hemispheric lateralization ranging (−0.019 0.039] for pars opercularis and (−0.001 0.04] for pars triangularis stimulation (Figure 4). For conduction speed (*v* = 10 – 19 *m/s*) the *LI*’s were positive while for low conduction speed (*v* = 5 - 10 *m/s*), *LI*’s were negative.

**Figure 4:**
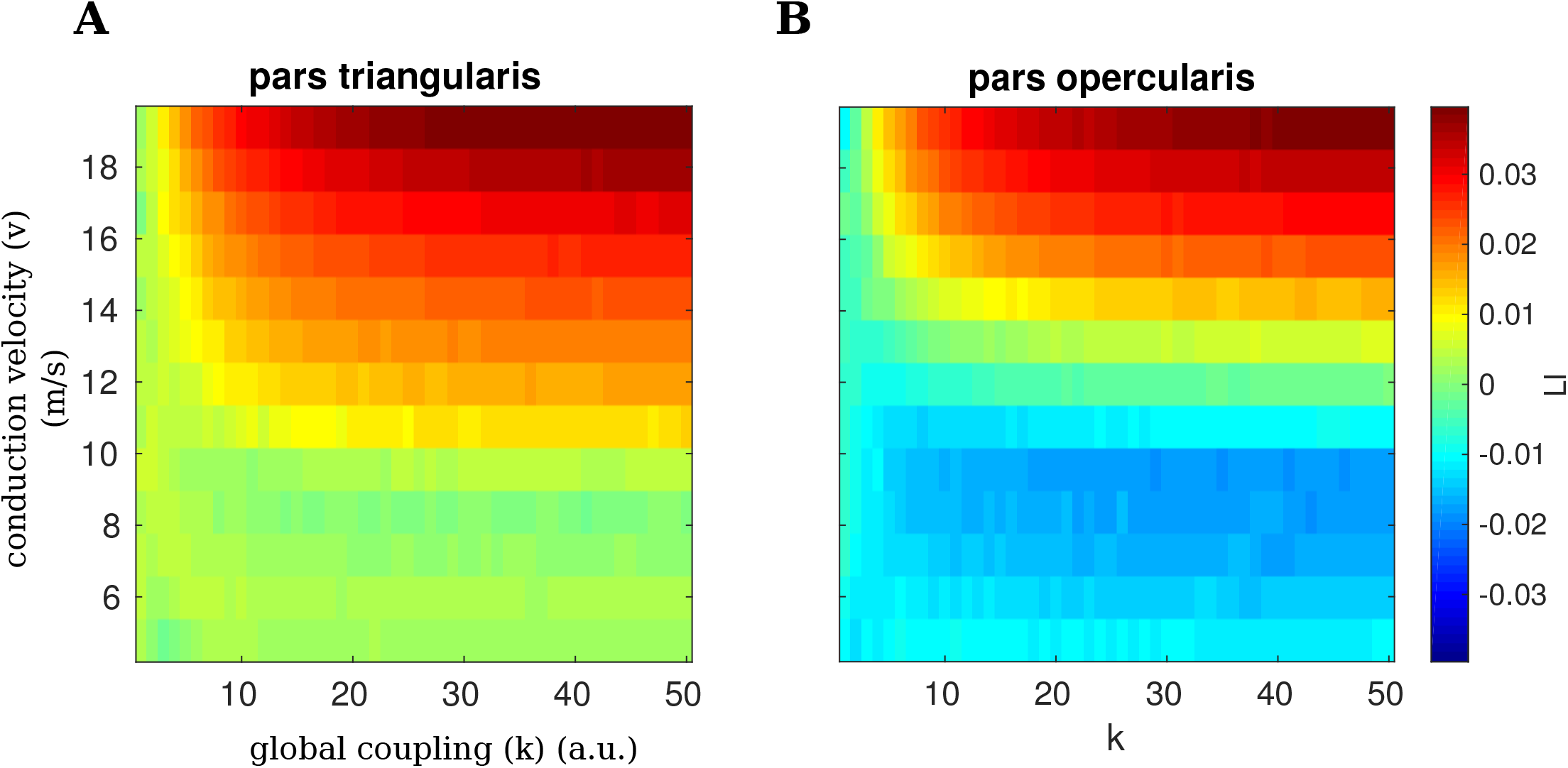
Exploring lateralization effects in language networks: *LI* of 40 Hz spectral power under the parametric variation of global coupling (*k*) and conduction velocity (*v*) in the large-scale neural model while setting seed regions of 40 Hz oscillation at sub-divisions of Broca’s area A) pars triangularis and B) pars opercularis.

Additionally, we tested for the scenarios of negative time-delay in system of equations (1–2) which can be numerically realized as situations where time-delayed contributions of phases of non-AC parcels θ_n_(t – τ_np_) is coupled to θ_A_(t) for defining the derivative of phase 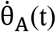. Physiologically this can be contextualized to a scenario where auditory cortical nodes are driven by feedback from other brain regions rather than feed-forward communication captured by positive τ_np_. Numerical integration of the revised equation 1–2 did not show emergence of 40 Hz oscillation. This result was unchanged whether auditory cortices or Broca’s area was chosen as areas with intrinsic frequency set to 40 Hz.

## Discussion

In the present study, we evaluated the role of whole-brain structural connectome comprising fairly detailed brain parcellation in shaping up the lateralization of brain dynamics. The primary hypothesis was validated using a canonical model for functional brain lateralization at two stages. First, we replicated the earlier findings of entrained 40 Hz oscillations triggered by amplitude modulation of the auditory stimuli, corroborating a number of previous studies (Galambos et al., 1981; Hari et al., 1989; Linden et al., 1987). While earlier studies concentrated predominantly on right hemispheric dominance of primary auditory areas (Ross et al., 2005), our results additionally demonstrate that the generators of auditory steady-state responses (ASSRs) need not be restricted to primary auditory areas only, instead of distributed over the entire right hemisphere suggesting a role of distributed network assemblies in processing sensory information. Importantly, we also demonstrate the lateralization of ASSR, although reported mostly in group-level studies (e.g., Ross et al., 2005), can also be observed at individual participant level in 65% (binaural), 60% (monoaural left), 70% (monoaural right) of the sample studied. Finally, by simulating auditory nodes as externally driven functional units in a whole-brain network of Kuramoto oscillators (Kuramoto, 1984), we could generate the functional lateralization of ASSR. Second, when exchanging identities of areas that receive direct environmental input, e.g., replacing auditory areas that receive tonal cues with Broca’s area receiving speech signals, we could predict the left hemispheric dominance of functional brain responses as reported by several language studies (Szaflarski et al., 2006; Riès et al., 2016; Olulade et al., 2020). Interestingly, our model also propose time-delays in coupling can affect the lateralization, e.g., allow transitions of left to right lateralization (Figure 4), which can provide important insights on why lateralization shifts are observed during ageing (Olulade et al., 2020) that is often associated with neurophysiological time-delays. Thus, two levels of validation – prediction of right hemispheric dominance during generation of ASSR and left hemispheric dominance of language qualifies this model as a canonical model for brain lateralization, which remains elusive to date to the best of our knowledge.

### Sources beyond the auditory cortex for generation of ASSR

eLORETA was used to locate 40 Hz ASSRs sources during binaural and both monaural conditions, as it has been shown to have significant control over the false-positive ratio in the distributed dipole condition (Halder et al., 2019). Subsequently, strong activation beyond the primary auditory cortex is reported in the present study. For instance, brain regions in the inferior parietal gyrus, pre-central and post-central gyrus, inferior and middle frontal gyrus, and occipital cortex (Figure 2-1) are in line with earlier findings on reconstructed 40 Hz ASSRs sources with equivalent dipole modeling. Earlier research identified distributed sources in both cortical and subcortical regions during 4, 20, 40, and 80 Hz ASSRs (Farahani et al., 2017). Here, we demonstrate right hemispheric dominance during binaural condition extends beyond primary auditory areas and can contribute to right hemispheric lateralization (Figure 2-1, Figure 2). During every stimulation condition, prominent sources were found among bilateral supra-temporal gyri, pre-central and post-central gyri. Activation in superior temporal gyrus (STG) corroborates with earlier findings (Mäkelä et al., 1987). Several sources in spatially distinct brain regions may also reflect different stages information processing hierarchy during binaural and both monaural conditions. In a PET-weighted LORETA neuroimaging study Reyes and colleagues reported prominent activation in the right temporal lobe and right parietal lobe along with activations in the right frontal lobe during the monaural right condition (Reyes et al., 2005). Several studies have also reported anatomical projections from STG to the frontal cortex (Hackett, 2011; Kaas et al., 2000; Plakke et al., 2014). Wang and colleagues identified a functional network comprising of the frontal cortex and superior temporal regions that are sensitive to tone repetition pattern, associated with human’s unique ability for language processing (Wang et al., 2015). Hence, the results presented here support the emerging view that auditory processing at the sensory level requires other brain areas beyond primary auditory cortices.

### Generative mechanisms of asymmetric lateralization of functional brain responses

ASSRs involve synchronizing distributed neuronal assemblies to periodic external input (Pastor et al., 2002; Reyes et al., 2005). Heschl’s gyri situated in primary auditory cortices are known to be the first cortical structure to receive auditory information. Subsequently, information from primary auditory cortices is segregated to the specialized higher-order cortical networks to resolve and process features of auditory stimuli. Mišić and colleagues suggested that asymmetry in communication pathways among both primary auditory cortices to other brain regions may contribute to functional lateralization in the auditory networks (Mišić et al., 2018). However, the generative mechanisms of such emergent functional lateralization remained elusive from previous studies. Using a neurodynamical whole-brain model constrained by biophysically realistic structural parameters, we capture the right hemispheric dominance of entrainment – a resonant physiological phenomenon. A more explicit validation of this hypothesis came from our source-level analysis, which revealed almost all the top sources showed right hemispheric dominance in terms of the number of sources and their power during binaural stimulation.

Another key finding of our study is that the right hemisphere at the source level was mostly dominant during every stimulation condition. However, if we compare activations in the left hemisphere among conditions, we found that the left auditory cortex has many sources during the monaural left condition than binaural and monaural right conditions. Specifically, left middle temporal gyri are activated in the monaural left condition but not during the monaural right. This result implies greater activation during ipsilateral ear stimulation compared to the contralateral ear stimulation. In an EEG study, Reyes and colleagues (Reyes et al., 2005) reported ipsilateral dominance in the temporal lobe /inferior parietal lobe (IPL) during 40 Hz - ASSRs. The dominance of ipsilateral (right) PoG and IPL during the monaural right condition was also reported by Pastor and Reyes, respectively, during 40 Hz ASSRs (Pastor et al., 2002; Reyes et al., 2005).

Using the Broca’s area as trigger zones of entrainment, we could simulate the scenario left-hemispheric dominance often observed with language-related brain responses. Our simulation results suggest possible sources of heterogeneous laterality indices in Broca’s area’s subdivisions – pars triangularis and pars opercularis. Given these findings, one could further investigate the other major sub-divisions in the brain with high-resolution brain parcellation schemes to address the effects of compensatory mechanisms during stroke recovery or other neurological conditions. While left hemispheric dominance for language processing is considered sacrosanct (Olulade et al., 2020), compensating right-hemispheric analog to Broca’s area is associated with recovery from stroke (Xing et al., 2016) can be understood with our model by making appropriate changes in the parameter space.

## Limitations and Conclusion

While pondering over these results, we should also be mindful of certain obvious caveats for contextualizing our results towards formulating a general theory of functional brain lateralization. Although we did not test the participants on their linguistic or auditory processing skills, all our participants were self-declared bi-literates and, in some cases, knew more than two Indian languages. Another critical point to note is that we did not choose participants based on their musical training, which can be a potential future direction for pattern differences. Thus, an exciting extension for the asymmetrical differences we report, can be explored in data sets present in the literature from neurodevelopmental disorders such as autism and neuropsychiatric disorders like schizophrenia. An important direction of ASSRs research is to define normative patterns of cortical auditory processing beyond simple audiological tests. We propose that our results be the most valuable, and more focused research delineating our neural disorders patterns is needed in the future. Another significant limitation of our model is that we did not consider subcortical areas in the network interactions. While certainly, this is an approach we want to take in the future, the current source localization methods become unreliable for deeper sources such as the thalamus and brain stem.

In summary, characterizing hemispheric dominance in the functional specialization of sensory processing is a fundamental question in cognitive neuroscience. Many factors may be involved to influence lateralization in brain, including anatomical asymmetries, stimulus designs, gender and handedness of the participant (Hutsler et al., 2003; Melynyte et al., 2017; Tervaniemi et al., 2000). Nonetheless, we have demonstrated that the structural connectome gives rise to two crucial physiological constraints: time-delays of propagation of information among brain areas and possible neural covariates dependent on tissue properties such as white matter density myelination that are also subject to neuroplasticity. In a nutshell, the time delay and global network coupling can be linked to two alternative plasticity mechanisms – experience/ age-dependent myelination/ demyelination and consolidation of between-network synaptic weights. Using our approach, both the aforementioned mechanisms can be studied to understand neuroplastic changes in brain responses in volunteers’ groups, such as musicians whose auditory systems may have higher fidelity and across life span aging cohorts where the brain’s tissue parameters have changed. Our model’s validation in such data sets will undoubtedly be one of our main targets in the future.

## Data and code Accessibility

The raw data have not been deposited on a public repository due to ethical considerations and Institution guidelines. Anonymized and processed EEG data, and codes to generate figures are uploaded at https://osf.io/sftqj/?view_only=76c7722b4b12465097246d2b4e80ce32.

## Author contributions

NK - Formal analysis, investigation, methodology, software, visualization, validation and writing original draft.

AJ - investigation and data curation.

DR - supervision, methodology, writing-reviewing and editing and funding acquisition.

AB - conceptualization, supervision, data curation, methodology, investigation, formal analysis, writing-reviewing and editing and funding acquisition.

## Acknowledgments

The study was supported by NBRC Core funds. DR was supported by the Ramalingaswami fellowship (BT/RLF/Re-entry/07/2014) from DBT and extramural grant SR/CSRI/21/2016/ from Department of Science and Technology (DST), Ministry of Science and Technology, Government of India. AB is supported by grant number F. NO.K-15015/42/2018/SP-V from Ministry of Youth Affairs and Sports, Government of India. AB and DR are jointly supported by BT/MED-III/NBRC/Flagship/Program/2019 from DBT. We sincerely thank late Dr. Jeffrey Michael Valla for constructive comments on an earlier version of this manuscript and Dr. Subham Kumar for helpful discussions and suggestions. Amit Kumar Jaiswal is currently placed as a Scientist at Elekta, Inc and as a PhD student at Aalto university, Finland.

## Extended data figure legends

**Figure 1-1:**
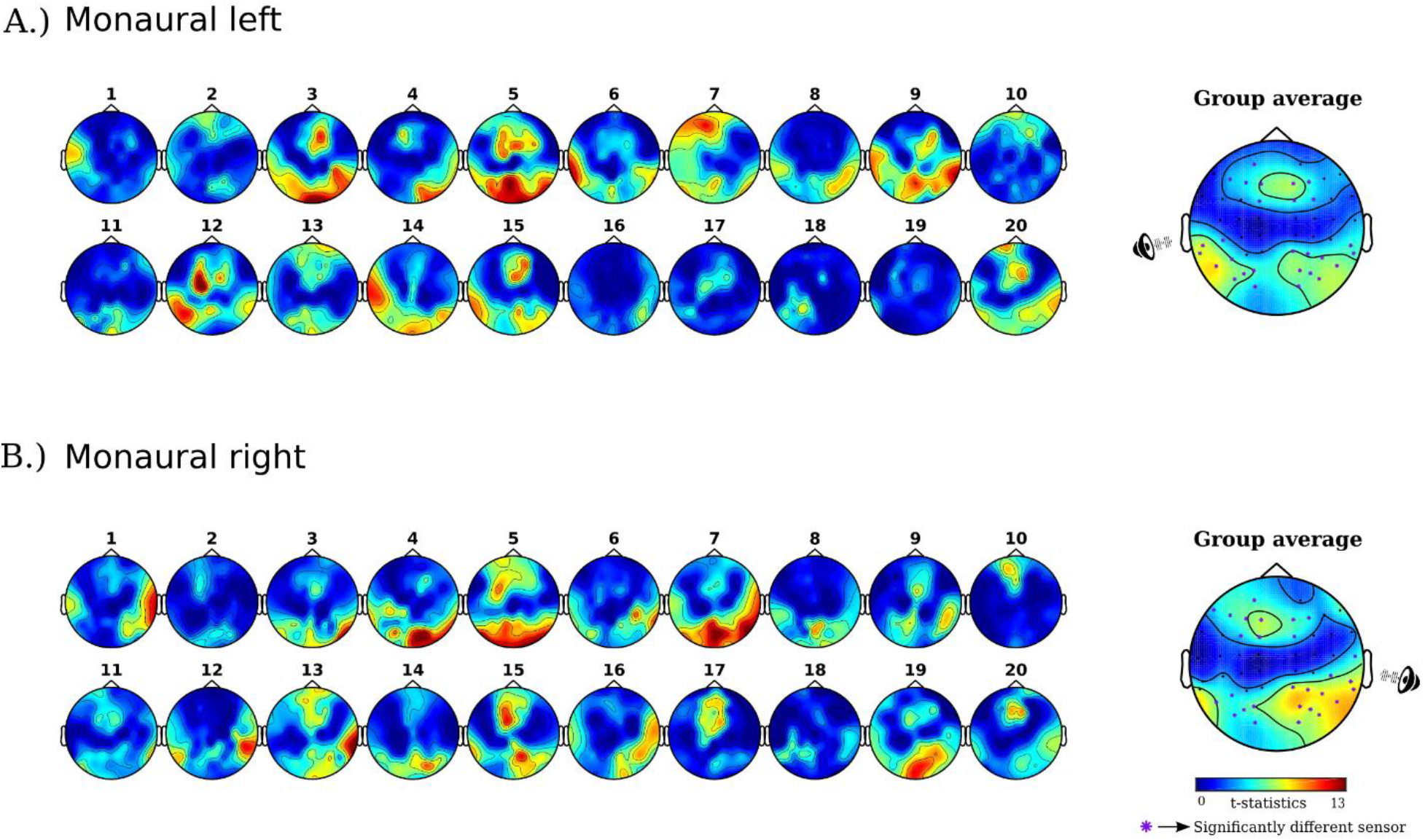
Subject-wise and group brain response during the monaural conditions: Topographical distribution of 40 Hz spectral power during the presentation of periodic auditory stimuli at 40 Hz during (A) monaural left and (B) monaural right condition.

**Figure 1-2:**
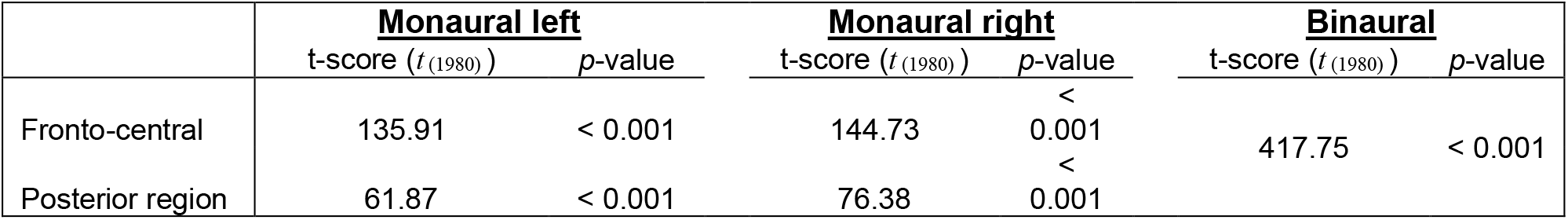
t-statistic and *p*-values of significant 40 Hz ASSRs clusters over the scalp locations for monaural and binaural stimulation.

**Figure 2-1:**
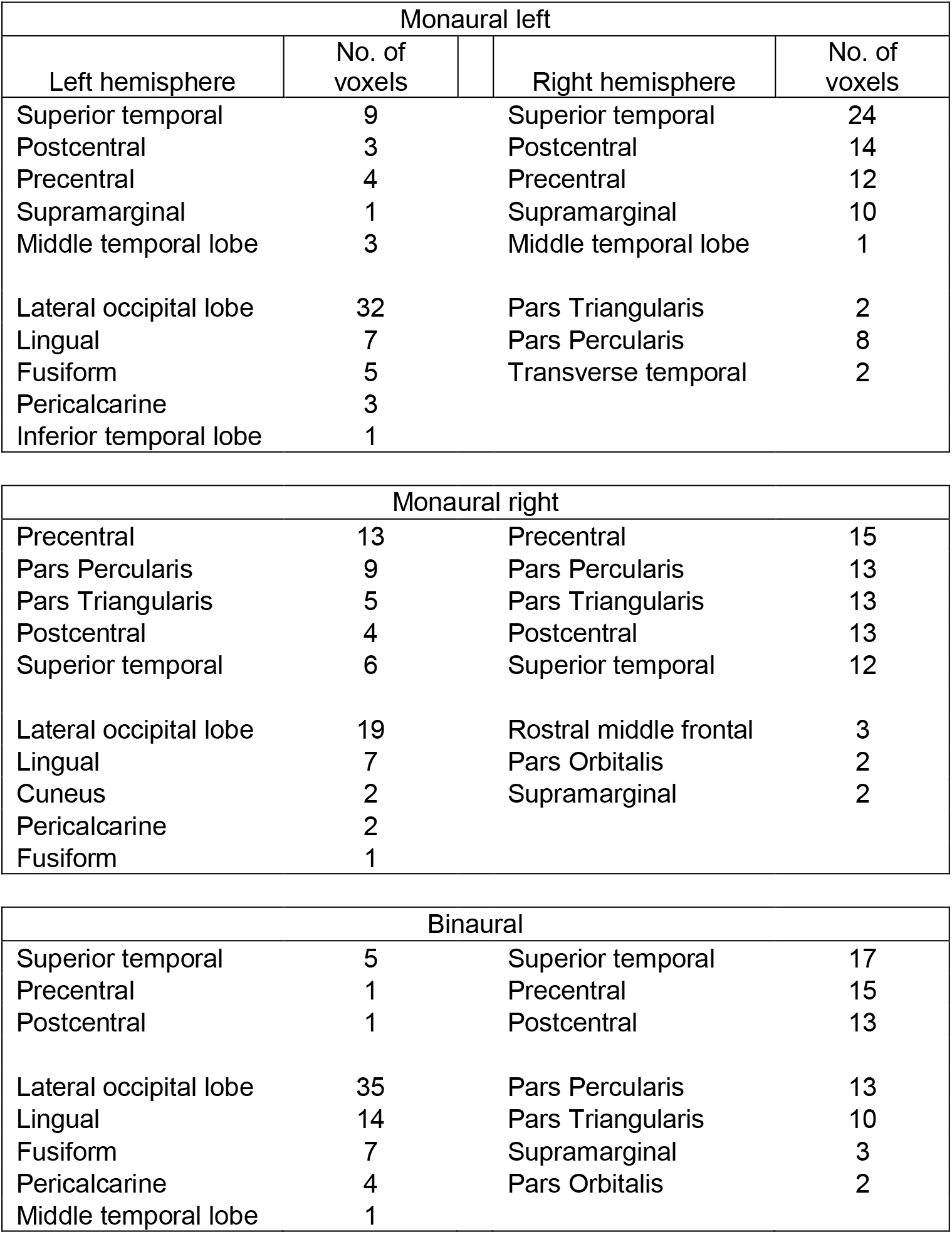
List of 40 Hz ASSRs source labels with the number of voxels activated in the left and right hemispheres.

## Footnotes

1 A subset of this data (10 volunteers) were used to validate the applicability of source localization methods previously: Halder, T; Talwar, S., Jaiswal, A.K., Banerjee, A.(2019): Quantitative evaluation in estimating sources underlying brain oscillations using current source density methods and beamformer approaches. eNeuro. 2019 Jul-Aug; 6(4): ENEURO.0170-19.2019.

